# Bioluminescence imaging of *Cyp1a1-*luciferase reporter mice demonstrates prolonged activation of the aryl hydrocarbon receptor in the lung

**DOI:** 10.1101/2023.05.30.542862

**Authors:** Nicolas Veland, Hannah J Gleneadie, Karen E Brown, Alessandro Sardini, Joaquim Pombo, Andrew Dimond, Vanessa Burns, Karen Sarkisyan, Chris Schiering, Zoe Webster, Matthias Merkenschlager, Amanda G Fisher

## Abstract

Aryl hydrocarbon receptor (AHR) signalling integrates biological processes that sense and respond to environmental, dietary, and metabolic challenges to ensure tissue homeostasis. AHR is a transcription factor that is inactive in the cytosol but upon encounter with ligand translocates to the nucleus and drives the expression of AHR targets, including genes of the cytochrome P4501 family of enzymes such as *Cyp1a1*. To dynamically visualise AHR activity *in vivo,* we generated reporter mice in which firefly luciferase (*Fluc*) was non-disruptively targeted into the endogenous *Cyp1a1* locus. Exposure of these animals to FICZ, 3-MC or to dietary I3C induced strong bioluminescence signal and *Cyp1a1* expression in many organs including liver, lung and intestine. Longitudinal studies revealed that AHR activity was surprisingly long-lived in the lung, with sustained *Cyp1a1* expression evident in discrete populations of cells including columnar epithelia around bronchioles. Our data link diet to lung physiology and also reveal the power of bespoke *Cyp1a1-Fluc* reporters to longitudinally monitor AHR activity *in vivo*.

## Introduction

The aryl hydrocarbon receptor regulates cellular physiology and organ homeostasis ^1, 2^. It was identified in the early 1990s as an environmental-sensor, with structural similarity to the class 1 basic helix-loop-helix-PER-ARNT-SIM (bHLH-PAS) family of transcription factors ^3–5^ and subsequently shown to be activated by a range of ligands ^6^. AHR recognises external xenobiotics, such as the polycyclic aromatic hydrocarbon dioxin, as well as endogenous metabolites including a plethora of compounds derived from tryptophan and dietary components generated by microbiota and host metabolism ^1, 7–10^. AHR is maintained in an inactive state in the cytoplasm, supported by a chaperone complex that includes 90 kDa heat shock protein (HSP90), AHR-interacting protein (AIP), co-chaperone p23 and SRC protein kinase. Ligand binding causes AIP to dissociate and triggers conformational changes that lead to the import of the complex into the nucleus where AHR binds to AHR nuclear translocator (ARNT, also known as HIF1β) and drives the expression of multiple target genes ^1^. Importantly, AHR activity induces the expression of enzymes of the cytochrome P450 family (*Cyp1a1, Cyp1b1*) which are capable of oxygenating and metabolically degrading endogenous and exogenous high affinity ligands ^11–16^. In addition, AHR activity induces expression of the AHR repressor (AHRR), which shares homology with AHR, ARNT and TiParp ^17, 18^ and competes with the AHR-ligand complex for ARNT binding, thereby creating a negative feedback loop that regulates AHR activation. Finally, AHR also regulates the expression of TiParp which in turn mediates the ribosylation and degradation of AHR ^19^. In this setting, interactions between AHR and ligand stimulate *Cyp1a1*, *Ahrr* and *TiParp* expression that subsequently act to degrade AHR ligand, reduce AHR availability, and counter AHR activation. Failures in this self-limiting process that lead to a dysregulated AHR pathway are linked to disease pathology and increased cancer risk ^20–36^.

Our appreciation of the importance of AHR signalling in sensing environmental and pathogen exposures, regulating tissue physiology, immune responses, and disease ontogeny, has increased substantially over the last decade. In particular, advances in the metabolic profiling of dietary response ^37, 38^, single-cell transcriptomic analysis of complex tissues ^30, 39, 40^, and assessing both canonical and alterative AHR ligands ^41^ have bolstered knowledge of the pleiotropic roles AHR signalling can play *in vivo.* Despite this, reagents that enable AHR activity to be reliably monitored in living tissues remain surprisingly limited. For example, although several models are available to examine the impact of deleting *Ahr* or *Ahr*-associated genes in cells, tissues and animals ^42–44^, routine cellular tracking of AHR/AHR-associated proteins using conventional antibody-based flow cytometry has remained elusive. Instead, endogenous tagging of AHR and AHR-associated proteins with fluorophores or other molecular adapters has been used to visualise these proteins in experimental settings *in vitro* or *ex vivo* ^45^.

Since *Cyp1a1* expression is dependent on AHR activation ^13–15^, induction of *Cyp1a1* is a useful surrogate for AHR activity. On this basis, several prior studies generated transgenic mouse lines that contained *Cyp1a1* promoter sequences, derived from rat or human, cloned upstream of reporter genes such as *CAT, luciferase* or *GFP* ^46–49^, reviewed in ^44^. Such reporters have provided invaluable tools for assessing *Cyp1a1* responses to different environmental stimulants, but may not contain the full repertoire of genetic regulatory elements available within the endogenous *Cyp1a1* locus, which normally serve to control expression in different cell types and developmental stages. To address this gap, and moreover, to develop robust murine reporters that enable endogenous AHR activity to be longitudinally and non-invasively imaged, we inserted firefly luciferase (*Fluc*) into the 3’UTR of the mouse *Cyp1a1* locus. Analogous approaches had previously been used by our group to successfully derive mouse embryonic stem cells (ESCs) and animal models in which the allelic expression of imprinted genes can be visualised throughout lifespan and across generations ^50–52^ or that allow dystrophin and utrophin gene expression to be simultaneously imaged throughout mouse development ^53^.

Here we describe the generation and properties of a bespoke *Cyp1a1-Fluc* knock-in mouse reporter that was designed to sensitively monitor AHR activity across murine life course. We show that *Cyp1a1* expression remains low during foetal development, but is inducible upon exposure to AHR ligands. In adults, *in vivo* challenge with the high affinity endogenous ligand 6-formylindolo[3,2-β]carbazole (FICZ), or the environmental pollutant and AHR agonist 3- methylcholanthrene (3-MC), results in strong *Cyp1a1*-derived bioluminescence signal in intestine, lung, liver and heart tissues. We show that dietary exposure to indole 3-carbinol (I3C) also provokes durable *Cyp1a1* expression within the gastrointestinal track and among discrete populations of epithelial, endothelial and smooth muscle cells that are resident in the adult lung.

## Results

### A luciferase-based endogenous *Cyp1a1* reporter that monitors AHR activity *in vivo*

To generate a reporter for *Cyp1a1-*expression in pluripotent mouse ESCs, firefly luciferase (*Fluc*) was inserted into the 3’UTR of the endogenous *Cyp1a1* locus, downstream of exon 7 (Figure 1a summarises the targeting strategy). Self-cleaving T2A sites ensure that Cyp1a1 and luciferase polypeptides are generated from a single *Cyp1a1-Fluc* mRNA transcript, while preserving the function of the targeted allele ^54, 55^. Using this approach, two heterozygous *Cyp1a1^F^*^+/-^ ESC clones were generated, 1B2 and 1D10, which were verified by DNA sequencing. Treatment of either clone with FICZ for 5 hours resulted in significant bioluminescence signal (blue-green) upon addition of D-luciferin (Figure 1b, and quantified in bar chart, right). Consistent with this we detected significant increases in *Cyp1a1* mRNA following FICZ exposure, as compared to vehicle treated controls (Figure 1c). As anticipated, control wild type ESCs (WT) showed a similar increase in *Cyp1a1* expression in response to FICZ treatment (Figure 1c), without detectable bioluminescence signal (Figure 1b, WT). Exposure of 1B2 and 1D10 clones to 3-MC also provoked a significant increase in *Cyp1a1* mRNA detection relative to vehicle controls (Figures S1a). These data are consistent with increased *Cyp1a1* expression and luciferase activity in targeted ESC clones following exposure to AHR ligands. Clone 1B2 was then used to create mouse lines where AHR-ligand responses could be investigated in a whole organism setting.

**Figure 1.**
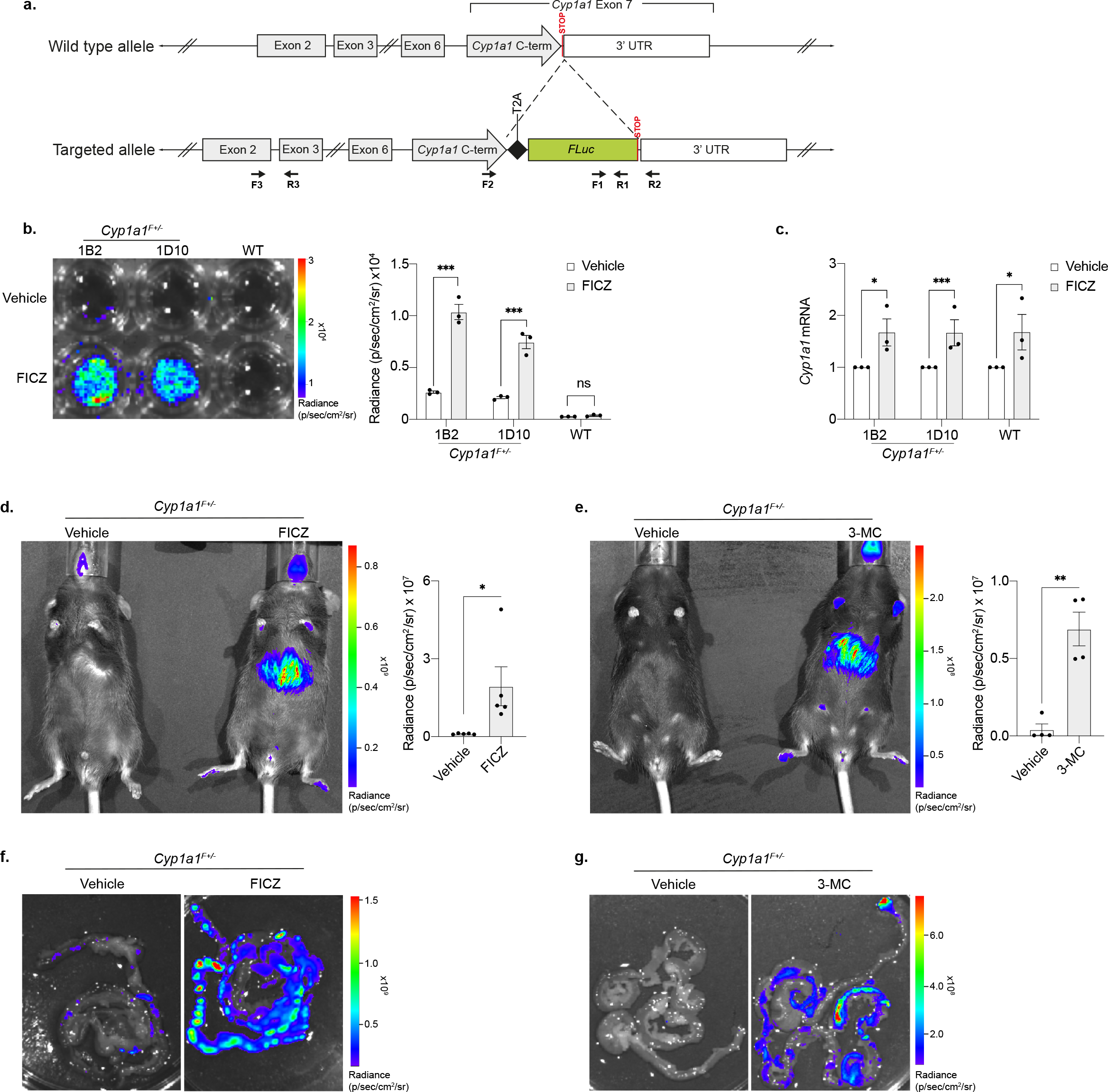
Generating a luciferase-based allelic reporter of endogenous *Cyp1a1* expression. **a.** Diagram of the gene targeting strategy used to generate knock in *Cyp1a1^F^* reporter mESCs and mouse lines. A firefly luciferase (*Fluc*) gene was inserted just before the stop codon in exon 7 of endogenous *Cyp1a1,* and it was separated from the C-terminal region by a T2A sequence. Arrows indicate PCR primers: F1 and R1 were used for firefly luciferase (*Fluc*) genotyping (see S1b), F2 and R2 for *Cyp1a1* wild type allele genotyping (see S1b), and F3 and R3 for mRNA quantification. **b.** Representative bioluminescence image (left) and flux quantification (right) of two *Cyp1a1^F^* mESC clones (1B2 and 1D10) shown alongside the parental wild type (WT) mESC line after 4-hour exposure to FICZ or vehicle. Bars show the mean of 3 replicates +/- SEM, with paired t-tests to compare vehicle with FICZ treated samples for each cell line. **c.** RT-qPCR of *Cyp1a1* mRNA expression from *Cyp1a1^F^* (1B2 and 1D10) and WT mESCs following 4-hour FICZ treatment. Levels of *Cyp1a1* mRNA are normalised to *Gapdh* mRNA and shown relative to the vehicle control. Bars show mean (n=3) +/- SEM with paired t-tests to compare vehicle with FICZ treated samples. **d.** Bioluminescence imaging of heterozygote (*Cyp1a1^F+/-^*) adult mice 5 hours post intraperitoneal (IP) injection with vehicle or FICZ. Representative image (left) and whole-body quantification (right). Bars show mean (n=5) +/- SEM with an unpaired t-test to compare vehicle with FICZ treated sample. **e.** Representative image (left) and whole-body quantification (right) of *Cyp1a1^F+/-^* adult mice 5 hours post IP injection with vehicle or 3-MC. Bars show mean (n=4) +/- SEM with an unpaired t-test to compare vehicle and 3-MC treated samples. **f** and **g.** Representative bioluminescence images (n=3) of intestine dissected from *Cyp1a1^F+/-^* mice 5 hours post FICZ (**f**) or 3-MC (**g**) injection. *p<0.05, **p<0.01, ***p<0.001.

*Cyp1a1-Fluc* knock-in animals were derived, and genotyped as described in Figure S1b. Whole body bioluminescence imaging of these mice revealed *Cyp1a1*-derived flux signal in living (anaesthetised) heterozygous *Cyp1a1^F+/-^* animals 5 hours after injection with FIZC or 3-MC (Figures 1d and 1e respectively, compare with *Cyp1a1^F+/-^* animals injected with vehicle alone). Bioluminescence was detected in multiple tissues and was also verified posthumously, following dissection. For example, elevated bioluminescence signals throughout the gastrointestinal tracts of FICZ or 3-MC treated animals were detected (Figures 1f and 1g, respectively), as compared to vehicle treated *Cyp1a1^F+/-^* controls. These data show that exposure to AHR ligands *in vivo* induces the expression of *Cyp1a1*-derived luciferase in reporter mice, that can be visualised and quantified by bioluminescent imaging.

### Durable *Cyp1a1-Fluc* expression *in vivo* following challenge with FICZ or 3-MC

To investigate the duration of AHR-ligand responses *in vivo*, we performed longitudinal imaging and molecular analyses to track *Cyp1a1-Fluc* expression over time. Heterozygous reporter mice were examined 5 hours and 6 days after FICZ or 3-MC challenge, as outlined in Figure 2a. In response to FICZ, whole body bioluminescence imaging detected a variable but significant increase in flux signal relative to vehicle alone controls, that declined by day 6 (Figure 2b shows representative images [left], and signal quantification [right]). In response to 3-MC, strong bioluminescent signal was evident at 5 hours (Figure 2c). Closer inspection of animals sacrificed at each timepoint revealed significant increases in both luciferase and *Cyp1a1* mRNA expression in liver and lung samples 5 hours after FICZ exposure (Figures 2d and 2e), with persistent signal/expression detected in the lung 6 days after exposure. In these samples we detected similar increases in *Cyp1b1* expression in response to FICZ (Figure S2a), while *Cyp1a2* showed transient upregulation only in the liver (Figure S2b), consistent with the reported tissue-associated expression of this gene ^56^. FICZ exposure also provoked *Cyp1a1- Fluc* upregulation in the heart (Figure 2e), although a corresponding signal was not immediately visible by bioluminescence imaging (Figure 2d), most likely because heart is enriched with blood and absorbance can mask the detection of emitted photons ^53^. Collectively these data show that AHR remains active in lung 6 days after FICZ exposure.

**Figure 2:**
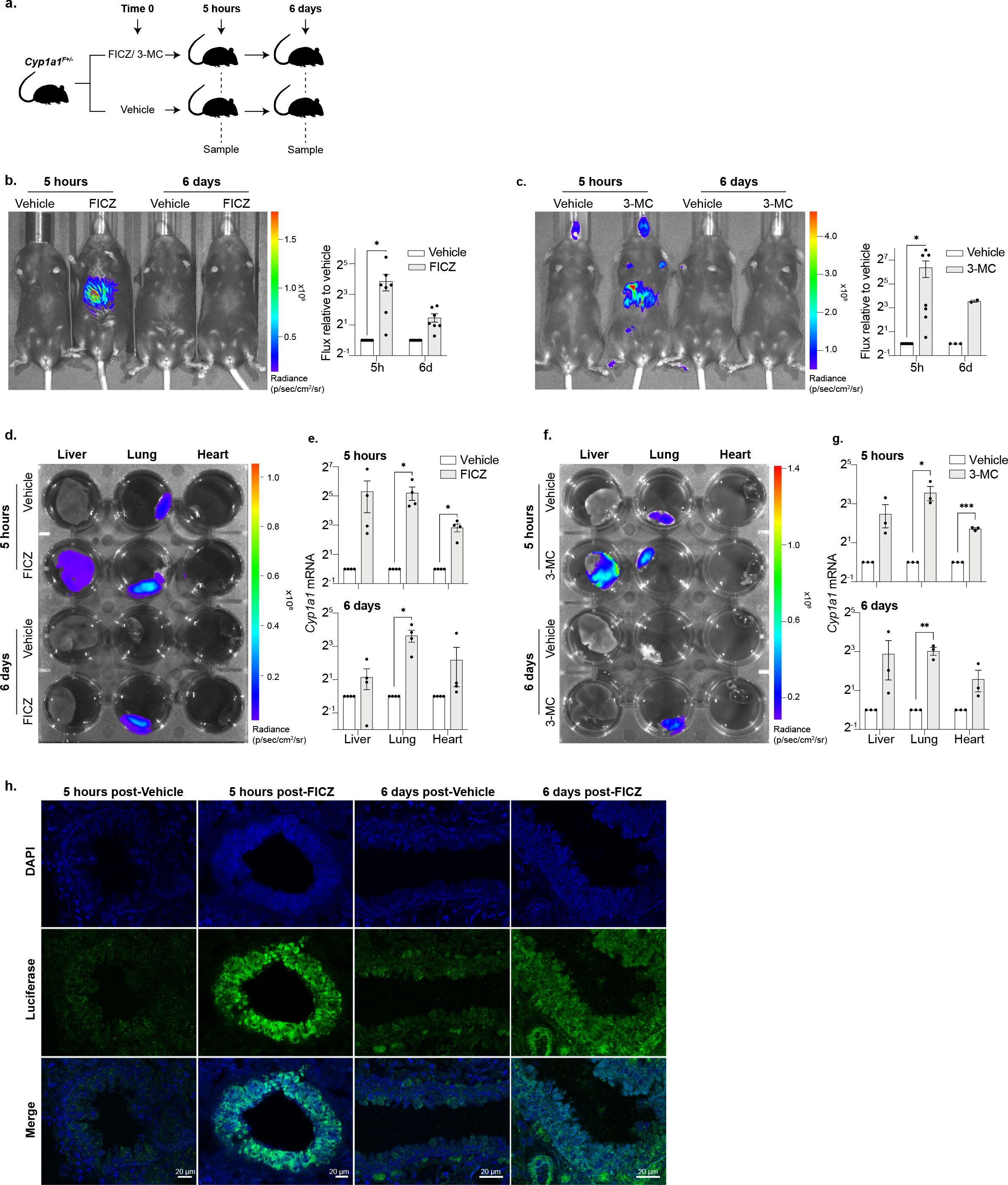
Longitudinal imaging of AHR activity following FICZ and 3-MC exposure *in vivo* **reveals prolonged *Cyp1a1* expression in the lung. a.** Experimental design diagram outlining the exposure duration and sampling protocol. *Cyp1a1^F+/-^* mice were IP injected with either an AHR ligand (FICZ or 3-MC) or vehicle (corn oil). Mice were sampled for *in vivo* and *ex vivo* bioluminescence imaging, RT-qPCR or immunofluorescence analysis at 5 hours and 6 days post ligand exposure. **b.** *In vivo* bioluminescence imaging of *Cyp1a1^F+/-^* adult mice 5 hours and 6 days post FICZ injection. Representative image (left) and whole-body quantification of radiance (right). Bars show mean flux relative to vehicle treated control +/- SEM. Unpaired t-tests were used to compare vehicle with FICZ-treated mice for each time point (*p<0.05). **c.** As in **b** but following 3-MC injection. **d-e.** Liver, lung, and heart were dissected from adult *Cyp1a1^F+/-^* mice 5 hours and 6 days after FICZ injection. **d.** Representative bioluminescence image of tissues following 2 minutes incubation in D-luciferin. **e.** RT-qPCR for *Cyp1a1* expression at 5 hours (upper) and 6 days (lower) post FICZ injection. *Cyp1a1* mRNA is normalised against *18S* rRNA and *Tbp* mRNA and each sample is shown relative to its vehicle treated counterpart. Bars show mean +/- SEM and t-tests with Holm-Sidak multiple comparison testing were used to compare vehicle with FICZ treated samples (adjusted p-values are shown *p<0.05, ***p<0.001). **f-g.** As in **d-e** for 3- MC treated samples. **b, c, e, g.** Graphs are shown in Log2 scale. **h.** Anti-luciferase immunofluorescence staining of 20 micron sections of lung tissue collected 5 hours (first two panels) or 6 days (final two panels) post injection with FICZ or vehicle. Upper panel shows staining for nuclei alone (DAPI), middle shows anti-luciferase staining and lower panel shows the two stains merged.

*Cyp1a1^F+/-^* mice injected with 3-MC showed bioluminescent signal in isolated liver and lung (Figure 2f), with prominent and durable *Cyp1a1*-derived mRNA expression again detected in the lung (Figure 2g, 6 days). These results therefore show that although exposure to FICZ or 3-MC results provoke different kinetics of AHR activation and ligand clearance ^13, 57, 58^, *Cyp1a1* responses to both agents in the lung were surprisingly long-lived. In addition, we noted low-level expression of *Cyp1a1*-derived signal in control (vehicle-treated) lung tissue (Figures 2d and 2f, top row middle), which infers that the basal expression of *Cyp1a1* in adult mouse lung might be higher than in other tissues. To identify cells within lung that respond to AHR ligand, immunofluorescence labelling was performed using anti-luciferase antibody to label cells expressing *Cyp1a1-Fluc* in tissue sections. Columnar epithelial cells that surround bronchioles were intensely labelled with anti-luciferase antibody following exposure to FICZ (Figure 2h, 5 hours post-FICZ, green). Six days after exposure, although the level of *Cyp1a1*-driven luciferase labelling was reduced (Figure 2h, right panels) these were still clearly above the levels seen in vehicle-exposed control lung. Closer inspection revealed that in addition to bronchiole epithelial cells, other cell types in the lung tissue expressed *Cyp1a1*-derived luciferase following FICZ treatment, as illustrated in Figures S2c and S2d. This included discrete populations of smooth muscle cells and vascular endothelium, identified by co-labelling with anti-αSMA and anti-CD31, respectively (Figure S2d).

### Expression of *Cyp1a1* during mouse ontogeny

To explore when *Cyp1a1* is expressed during mouse development, we performed bioluminescence imaging and molecular analysis of embryos generated from mating *Cyp1a1- Fluc* heterozygote and wild type mice. In prior studies, using a transgenic mouse line containing 8.5 kb of the rat *Cyp1a1* promoter linked to *lac*Z ^59^, *Cyp1a1*-driven expression was reported in many tissues throughout stages E7-E14. Despite this expectation, we did not observe any generalised expression of *Cyp1a1-*derived signal in *Cyp1a1^F^*^+/-^ reporter embryos sampled from E10 to E14.5 (Figure 3a). We have previously shown that bioluminescence can be sensitively imaged in developing mouse embryos using a range of different luciferase reporter lines ^50, 51, 53^, which excludes that failure to detect signal was simply due to a technical limitation in embryo imaging. Furthermore, exposure of E14.5 *Cyp1a1^F^*^+/-^ whole embryos or dissected tissues to FICZ *ex vivo*, resulted in abundant *Cyp1a1*-derived flux signal detection (Figures 3a and 3b) and *Cyp1a1* mRNA expression in embryonic heart, lung, liver and intestine (Figure 3c), as compared to vehicle controls. Taken together, these results clearly demonstrate that while *Cyp1a1* expression is normally low in the developing embryo, it can be induced upon exposure to AHR ligands. Differences between the results reported herein and those that were published previously ^59^ might therefore reflect differences in pathogen load or commensal microbes resident within different mouse colonies.

**Figure 3:**
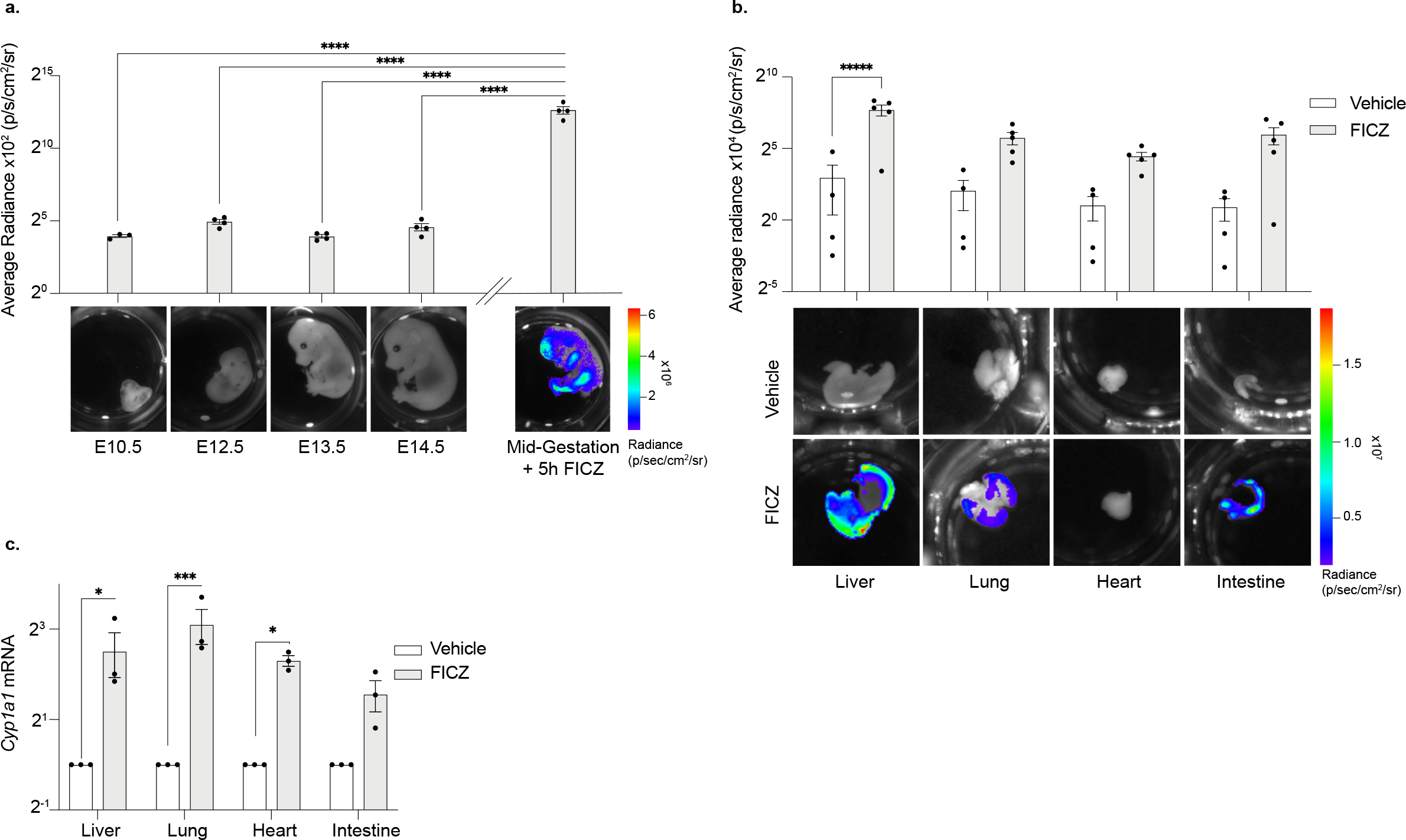
*Cyp1a1^F^* reporter expression during ontogeny. **a.** Bioluminescence images of *Cyp1a1^F+/-^* mouse embryos collected at E10.5, E12.5, E13.5 and E14.5 stages of gestation (without AHR stimulation). A mid-gestation embryo which was incubated with FICZ for 5 hours is shown alongside for comparison. Representative images are shown below and quantification above where bars show average radiance +/- SEM with a one-way ANOVA to compare the untreated embryos with the FICZ treated embryo (p<0.0001) with Dunnetts multiple comparison test. Adjusted p values are shown ****p<0.0001. **b.** Liver, lung, heart and intestine were dissected from E14.5 *Cyp1a1^F+/-^* embryos and incubated with vehicle or FICZ for 5 hours. *Ex vivo* bioluminescence images (upper) and quantification (lower) show increased luminescence in the liver, lung and intestine in response to FICZ. Bar graph shows average radiance +/- SEM with t-tests with Holm-Sidak multiple comparison testing were used to compare vehicle with FICZ treated samples (adjusted p-values are shown ****p<0.0001). **c.** RT-qPCR for *Cyp1a1* expression on samples from **b**. *Cyp1a1* mRNA is normalised against *18S* rRNA and *Tbp* mRNA and each sample is shown relative to its vehicle treated counterpart. Bars show mean +/- SEM with Holm-Sidak multiple comparison testing were used to compare vehicle with FICZ treated samples (adjusted p-values are shown * p<0.05, ***p<0.001) to compare vehicle with FICZ treated samples, (*p<0.05, ***p<0.001). **a-c** Graphs are shown on a Log2 scale.

### *Cyp1a1-Fluc* expression within the lung of reporter mice challenged with dietary I3C

Whereas mice that have been raised in a conventional setting are known to display *Cyp1a1* expression, for example in intestinal epithelial cells and in associated immune cells, those raised in germ-free conditions or exposed to lower levels of microbial factors express *Ahr*, *Ahrr* and *Cyp1a1* at lower levels ^60^. Exposure to I3C in diet, a natural product of glucobrassin hydrolysis, stimulates *Cyp1a1* activity in the intestine as well as in the liver ^27, 61^. Although I3C normally binds to AHR with low affinity, under acidic conditions I3C can be converted to indolo[3,2-β]carbazole, which has high affinity for AHR ^61^. We examined the impact of dietary I3C on *Cyp1a1* expression using *Cyp1a1^F^*^+/-^ and *Cyp1a1^F^*^+/+^ reporter mice. Animals were fed purified diet with or without I3C and imaged after 1 week (Figure 4a). To investigate the durability of I3C-induced *Cyp1a1* expression, mice that were exposed to control or I3C diet were then returned to normal chow for two further weeks before being imaged. As shown in Figure 4b, *Cyp1a1*-derived bioluminescence signal was readily detected in heterozygous and homozygous animals that had been fed I3C diet, with prominent signal evident in dissected lung samples (Figure S3). Molecular analysis across of a range of different tissues confirmed elevated *Cyp1a1* mRNA expression in the lung and colon of I3C exposed animals, compared with control diet samples (Figure 4c). Elevated bioluminescence signal was detected in the intestine of I3C-diet fed animals, as compared with control diet fed animals (Figure 4d), and signal intensity was noticeably higher in homozygous (*Cyp1a1^F^*^+/+^) than heterozygous (*Cyp1a1^F^*^+/-^) samples. Interestingly, although bioluminescent imaging showed luciferase activity throughout the intestine of I3C-fed *Cyp1a1^F^*^+/-^ animals, molecular analyses of *Cyp1a1*- mRNA in isolated regions of the gut (Figure 4c, right) detected significant increases only in the colon, rather than more proximal regions. This most likely reflects a known limitation of standard ‘bulk’ RNA analysis, where gene expression is averaged across a population of different cell types, and then normalised to standard ‘house-keeping’ genes. In such a setting, rarer cells expressing a gene of interest may be overlooked and remain undetected.

**Figure 4:**
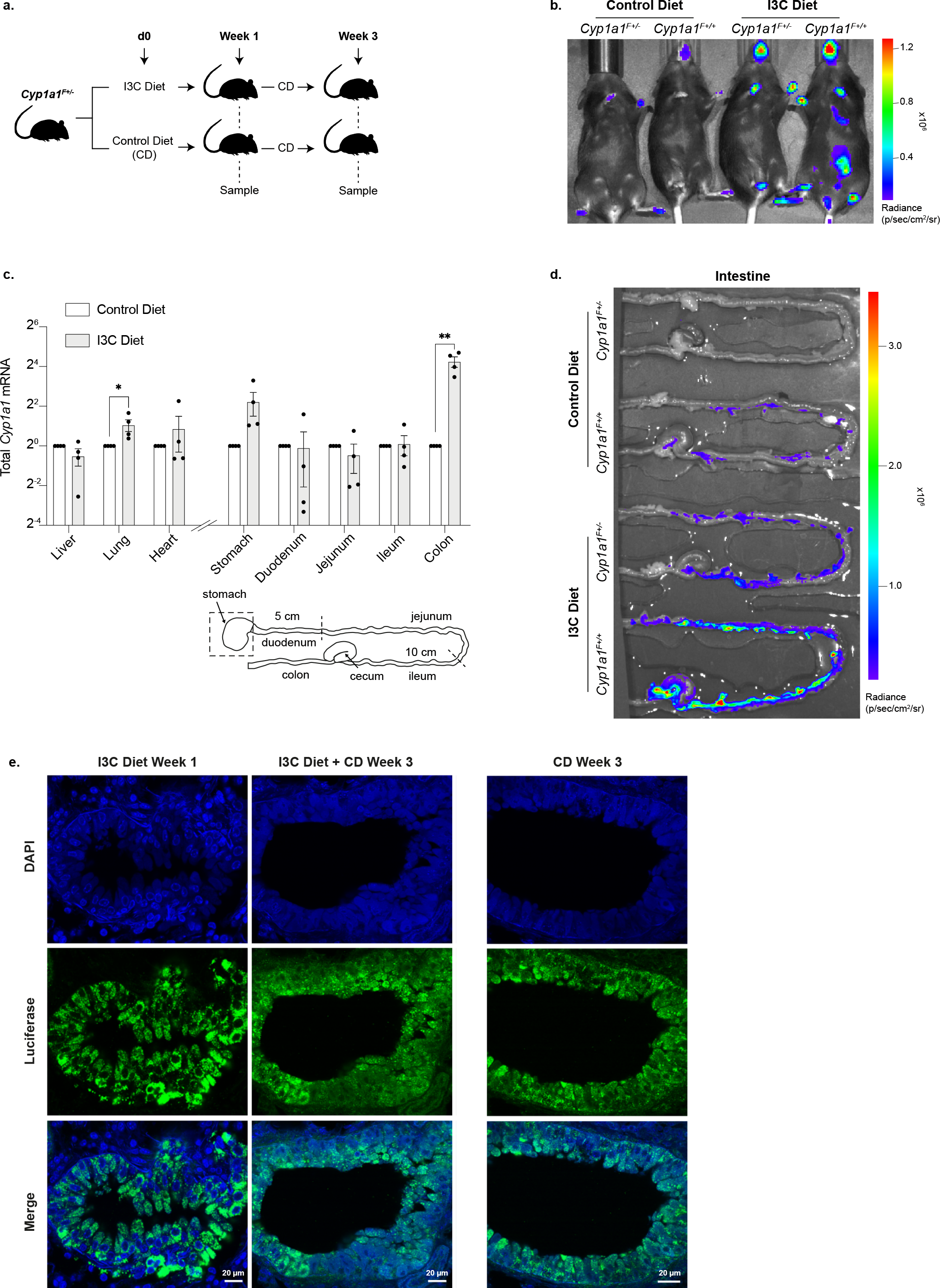
AHR activity in the intestine and lung of *Cyp1a1^F^* reporter mice fed with purified diet supplemented with I3C. **a.** Experimental design diagram outlining the exposure duration and sampling protocol. *Cyp1a1^F+/-^* adult mice were fed either a purified control diet (CD), or the same diet supplemented with I3C for one week. At this point mice were either sampled or switched to CD for a further two weeks and sampled at the three-week point. **b.** Representative bioluminescence image of *Cyp1a1^F+/-^* and *Cyp1a1^F+/+^* adult mice fed either purified control diet or I3C diet for one week. **c.** RT-qPCR on tissues samples isolated from adult mice following one week of purified control diet or I3C diet. *Cyp1a1* mRNA was normalised against *18S* rRNA and *Tbp* mRNA and shown relative to the corresponding control diet sample. Unpaired t-tests were used to compare control diet with I3C diet (*p<0.05, **p<0.01). A diagram of the different regions of the gut is shown below. **d.** Bioluminescence images of intestines dissected from *Cyp1a1^F+/-^* and *Cyp1a1^F+/+^* mice following one week of CD or I3C diet. Intestines are arranged corresponding to the gut diagram shown in figure **c**, with the stomach out of view of the image. **e.** Anti-luciferase immunofluorescence staining of lung tissue following one- week I3C diet (first panel); one-week I3C diet followed by two weeks CD (middle panel); or three weeks CD (final panel). * Mice housed in a non-SPF environment. Upper panels show nuclei staining with DAPI, middle panel shows luciferase staining and lower panel shows both channels merged.

To extend these findings and moreover to investigate *Cyp1a1* upregulation in the lung in response to dietary I3C, we performed immunofluorescence labelling using anti-luciferase antibody. We detected prominent labelling of columnar epithelium around lung bronchioles in mice fed I3C diet for a week (Figure 4e, left). Two weeks later, after being returned to a normal diet, appreciable *Cyp1a1*-driven luciferase expression remained in lung tissues (Figure 4e, centre). These data clearly showed that *Cyp1a1* expression by epithelial bronchioles in the lung was susceptible to dietary activation. However, we also noted that animals continuously fed purified control diet but housed in a non-SPF or conventional animal facility can also display elevated *Cyp1a1*-luciferase expression in these cell types (Figure 4e, right). Our results suggest that prolonged AHR activation in the lung can be stimulated by multiple agents and encompass different routes of exposure. Taken together, our data show how *Cyp1a1- Fluc* reporter mice can be used to identify sites of prolonged AHR activity *in vivo*, for example within the mouse lung, and thereby illustrate their utility as dynamic sensors of environmentally-induced AHR activation.

## Discussion

The aryl hydrocarbon receptor has fundamental roles in biology and AHR homologues are present in most animals from chordates to nematodes and molluscs ^62–65^. Evidence from vertebrates and invertebrates suggest that AHR signalling is an ancestral process that has, for example, underwritten the parallel development of sensory neural systems in both phyla. In vertebrates, AHR plays a crucial role in mediating responses to xenobiotics and in modulating adaptive immune responses to metabolites generated through bacterial, dietary and environmental exposures. This is best illustrated in the mammalian gastrointestinal tract where constant exposure to microbes and dietary ligands requires an epithelial barrier equipped with immune surveillance to protect and maintain health ^27, 28, 37, 60^. While the importance of AHR is widely appreciated, investigating the impacts of dietary exposures and the mechanisms that can resolve or potentiate AHR activity *in vivo* remains a challenge. Towards this goal we produced a bespoke mouse reporter line in which AHR-induced expression of endogenous *Cyp1a1* could be visualised longitudinally *in vivo* using bioluminescence. We predicted that this could offer two major advantages. First, because bioluminescence imaging does not require external excitation to generate signal, unlike conventional fluorophore-based approaches that monitor gene activity ^52^, we reasoned that this might improve signal detection by offering a high signal to noise ratio. Second, in contrast to previously generated *Cyp1a1*-promoter transgenic animals ^46, 48, 49, 59^, our strategy to create a ‘knock-in’ mouse by non-disruptive targeting of luciferase into the endogenous *Cyp1a1* locus should enable the normal dynamics of *Cyp1a1* expression to be accurately monitored. Our results show that *Cyp1a1^F+/-^* mice respond appropriately to AHR ligands such as FICZ and 3-MC, or dietary exposure to I3C, by upregulating luciferase expression. Increased *Cyp1a1- Fluc* expression was detected by bioluminescence imaging and confirmed by measuring luciferase mRNA in tissues using quantitative RT-qPCR, and protein distribution by immunofluorescence labelling with luciferase-specific antibody. *Cyp1a1-Fluc* adult mice housed in specific pathogen free conditions showed very low levels of luciferase reporter activity. Likewise, during foetal development we detected only minimal *Cyp1a1-Fluc* expression in embryos examined from E10.5 to E14.5. However, external exposure of mid-gestation embryos to FICZ (*ex vivo*) resulted in marked increases in *Cyp1a1-Fluc* expression, with bioluminescence signal evident in most major organs. These data support a view that AHR-signalling is inducible during mouse embryonic development ^59^, as well as in differentiating ESCs ^66^. Although prior studies with *Cyp1a1*-promoter transgenic mice have suggested that *Cyp1a1* expression is constitutive in embryos, with some evidence of temporal and spatial selectivity ^59^, in our hands *Cyp1a1* expression was uniformly low (basal) throughout ontogeny but remained inducible. While such differences could conceivably indicate the presence of negative regulatory elements at the endogenous *Cyp1a1* locus, it is perhaps more likely that such discrepancies merely reflect differences in the maternal availability of AHR ligands in animals housed under different conditions.

Inducible expression of *Cyp1a1* by alveolar and bronchiolar epithelial cells in response to smoking or hypoxia, has been described in humans and transgenic mice, respectively ^67, 68^. Here we show that *Cyp1a1* upregulation in bronchiolar epithelial cells was prolonged after exposure to either FICZ or to dietary I3C. The duration of *Cyp1a1* induction in these cells was longer than might be anticipated for ligands predicted to be susceptible to AHR-mediated metabolic degradation ^27, 69^. While the basis of this prolonged expression is not yet known, acute sensitivity of the respiratory system to altered AHR expression is well-documented ^70^ as is the role of AHR in modulating inflammatory lung diseases such as asthma, COPD and silicosis (reviewed in ^71^). Therefore, the provision of a bespoke *Cyp1a1* ‘knock-in’ reporter mouse line that accurately portrays the dynamics of AHR signalling longitudinally in individual animals will be of considerable value in evaluating the impacts and duration of repeated challenge.

Monitoring AHR responses *in vivo* is particularly difficult in complex tissues such as lung. Recent single-cell transcriptomic analysis of developing mouse lung reveals a diverse mixture of cell types ^72^, with eight different epithelial, six endothelial, and nine mesenchymal subtypes molecularly defined. Similar studies with human samples have confirmed this view, documenting a plethora of epithelial, endothelial, and mesenchymal cells that are integrated with immune cells to ensure airway development and function ^73–75^. Using our *Cyp1a1-Fluc* reporter mice we identified cell types within the lung that responded to FICZ or dietary challenge, including subsets of endothelium and smooth muscle. These observations align well with single cell RNA-seq data generated as part of the mouse cell atlas project ^76–78^ which showed *Cyp1a1* expression in three different subsets of lung endothelial cells, as well as myofibrogenic progenitors and smooth muscle. Our observation that dietary I3C exposure provokes prolonged activation of AHR in ‘barrier’ cell types in lung is important in understanding how encounter with respiratory pathogens may be affected by dietary or other environmental cues. It is well established that maternal diets enriched with AHR ligands can, for example, protect perinatal offspring from potentially lethal intestinal bacterial infection ^79^ as well as ameliorate colitis in adult mice (reviewed in ^1^). It is tempting to speculate that similar mechanisms operate in the lung where AHR signalling is known to afford anti-viral protection from agents such as Zika virus or SARS-CoV-2 ^80, 81^, as well as mediating reduced lung capacity through overt inflammation and increased mucin production ^82, 83^. Animal models that enable AHR activity to be quantitatively monitored through life in response to environmental changes, infection, disease and intervention, such as the *Cyp1a1-Fluc* reporter mice described here, offer important new tools to interrogate the fine balance between the therapeutic benefits and risks of modulating AHR activity *in vivo*.

## Acknowledgements

This work was funded by the Medical Research Council (MRC) (A.G.F. by MC_U120027516 and MC_UP_1605/12 and M.M by MC_UP_1605/11). N.V. received and ERDA award from the Institute of Clinical Sciences, Imperial College London. We would like to thank Ben Wiggins, Shwetha Raghunathan (NHLI, Imperial College London) and Mathew Van de Pette (MRC Toxicology Unit, Cambridge) for sharing their expertise and advising.

## Author contributions

N.V. and A.G.F conceptualised the study with input from C.S., K.S and M.M. The majority of experiments were performed by N.V, H.J.G., V.B and K.E.B, with expert help from A.S., Z.W., A.D. and J.P. A.G.F, N.V and H.J.G wrote the manuscript, with input from all co-authors.

## Methods

### Animal maintenance

All animal procedures were performed in accordance with the British Home Office Animal (Scientific Procedures) Act 1986. The mouse work was approved by the Imperial College AWERB committee and performed under a UK Home Office Project Licence and Personal Licences. Mice were housed in a SPF facility at temperatures of 21+/−2 °C; 45–65% humidity; 12-h light-dark cycle; with water and RM3 diet ad libitum. Tissues, wood blocks, and tunnels were used to enrich the environment. Experiments on adult mice were performed on animals between 3–16 weeks old.

### Generation of mESCs, mouse line and PCR genotyping

*Cyp1a1-Fluc* (referred to as *Cyp1a1^F^*) mESCs and mouse line were generated by OzGene, Australia. A firefly luciferase (*Fluc*) gene was inserted just before the stop codon in exon 7 of endogenous *Cyp1a1*, and it was separated from the C-terminal region by a T2A sequence (see Supplementary Figure 1a for details). Genotyping by PCR was carried out using HotStar Taq DNA Polymerase (Qiagen) according to manufacture conditions. Two independent PCR reactions were performed in parallel for each DNA sample. A first PCR reaction with two set of primer pairs at a final concentration of 0.2 μM each: one specific for *Fluc* and another specific for a region of wild type *CD79b* (Chr11: 17714036-17714620) that serves as internal control. A second PCR reaction with only one pair of primers at a final concentration of 0.4 μM specific for the wild type allele of *Cyp1a1.* Primer sequences are indicated in Table 1.

**Table 1.**
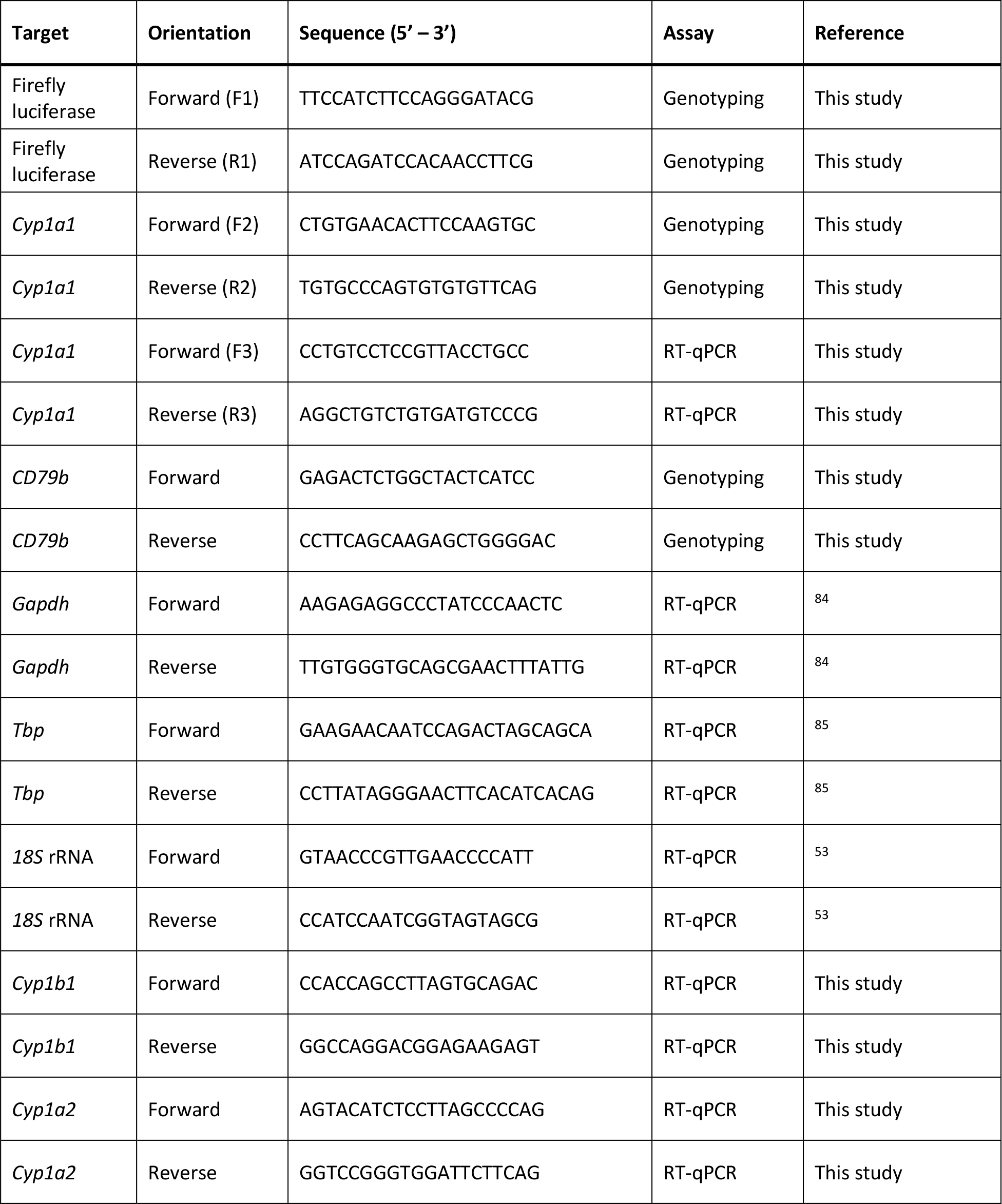
Primer sequences used in this study.

### mESC culture and treatment

C57BL/6 knock-in mESCs clones and Bruce4 parental wild-type mESCs were cultured on a layer of mitotically-inactivated mouse embryonic fibroblasts on 0.1% gelatin-coated dishes with KnockOut Dulbecco’s Modified Eagle’s Medium (Gibco), supplemented with 15% fetal bovine serum (Gibco), 0.5% penicillin-streptomycin (Gibco), 0.1 mM non-essential amino acids (Gibco), 2 mM L-glutamine (Gibco), 0.1 mM 2-mercaptoetanol (Sigma), 10^3^ U/mL of leukemia inhibitory factor (ESGRO, Millipore) and 2 μM of GSK-3 inhibitor IX (BIO, Selleckchem). Cells were incubated at 37° C with 5% CO2 and split every two to three days. Cells were treated with 10 nM FICZ (BML-GR206-0100, Enzo), 1 nM 3-MC (213942, Sigma) or DMSO (Sigma) as vehicle control.

### Animal studies

For *in vivo* experiments, adult mice were weight and intraperitoneal (IP) injected with FICZ (SML1489, Sigma) or 3-MC (213942, Sigma) freshly prepared in warm corn oil (Sigma) at 10mg/kg or 26.5mg/kg, respectively. Corn oil was used as vehicle control.

For timed mating, an adult male was set up with 2 adult females and morning plug checking was performed. The females were separated from males upon observation of vaginal plugs, at which point they were considered to be at E0.5 developmental stage.

For diet studies, adult mice were fed with purified diet E157453-047 (D12450J) or E157453- 047 (D12450J) containing 1000 mg/kg I3C (Sigma) provided by ssniff Spezialdiäten GmbH. Both diets were sterilized by gamma-irradiation at 25 kGy.

### *In vivo* and *ex vivo* bioluminescence imaging

To image mESCs, D-Luciferin (Perkin Elmer) was diluted in ESC medium to a final concentration of 150 μg/mL and added to mESCs 10 minutes prior to imaging. Cells were imaged using the IVIS Spectrum (Perkin Elmer) and Living Image software (version 4.3.1) to detect bioluminescence. All images were taken after 5 minutes exposure and at field of view (FOV) C with binning 4 and 0.5 depth using a stage temperature of 37° C.

For *in vivo* bioluminescence imaging experiments, adult mice were intraperitoneal (IP) injected with 0.15 mg/g D-Luciferin (Perkin Elmer), dissolved in dH2O. Mice were left conscious for 3 minutes to allow the D-Luciferin to circulate systemically and then anesthetized through isoflurane inhalation. At 10 minutes post-injection, mice were imaged using the IVIS Spectrum. Adult mice were imaged for 3 minutes using FOV C or D, binning 1 and 1.5 depth using a stage temperature of 37° C.

For *ex vivo* experiments, dissected tissues were incubated in 150 μg/mL D-Luciferin in DMEM medium without Phenol Red (Gibco) for 2 minutes prior to imaging for 3 minutes at FOV C or D with binning 1 and 0.5 depth using a stage temperature of 37° C.

For embryos imaging, pregnant females were first IP injected with 0.15 mg/g D-Luciferin and left conscious for 3 minutes to allow the D-Luciferin to circulate systemically, then mice were culled followed by embryo dissection. Embryos were placed in 24-well plates and incubated with freshly prepared 150 μg/mL D-Luciferin in DMEM/F12 medium (Gibco) for 2 minutes prior to imaging on the IVIS for 1 minute using FOV A, binning 4 and 0.75 depth.

Image analysis and bioluminescence quantification were carried out using the Living Image software (version 4.5.2) (Perkin Elmer). Briefly, regions of interest (ROI) were drawn around plate wells containing cells, tissues and embryos or around whole animals to calculate flux (p/s) and average radiance (p/s/cm^2^/sr) within the region.

### RNA extraction and RT-qPCR

RNA from mESC and tissue samples was purified using the RNeasy Mini Kit (Qiagen). Cells were lysed immediately after imaging with RLT buffer. Tissues were dissected and frozen in liquid nitrogen. Prior to RNA purification, frozen tissues were lysed in RLT buffer on the TissueLyser II (Qiagen) using 5 mm stainless steel beads (Qiagen) for 4 minutes at 24,000 rpm. Heart samples were incubated with 10 μg/ml Proteinase K at 55 °C for 1 hour. All tissue samples were then centrifuged at top speed for 3 minutes and total RNA was purified from the supernatant using the RNeasy Mini Kit (Qiagen) according to the manufacturer’s instructions, including on-column DNase digestion step using an RNase-Free DNase Set (Qiagen). After quantification, 2 μg of total RNA was used to perform cDNA synthesis with 10 μM random primers using the SuperScript III Reverse Transcriptase Kit (Invitrogen), following manufacturer’s instructions.

RT-qPCR was performed with 0.4 μM primers and using the QuantiTect SYBR Green PCR mix (Qiagen). Primer sequences are indicated in Table 1. Samples were analysed in 3 technical replicates. PCR reactions were carried out in a CFX thermocycler (Bio-Rad) for 40 cycles of a 2-step amplification protocol consisting of 94° C for 15 seconds and 60° C for 30 seconds. A disassociation final step to calculate melting temperature was included in all RT-qPCR experiments.

### Immunofluorescent microscopy on frozen tissue sections

Mouse lung or colon tissue was dissected, fixed in 10% Formalin solution (Sigma Aldrich), incubated with 30% sucrose solution for three days at 4°C and frozen in Optimal Cutting Temperature (OCT, Thermo) to form blocks. The tissue blocks were cryosectioned (20 µm) and mounted on microscope slides (Superfrost Plus Adhesion Microscope slides, VWR) and stored at -80°C. Thawed sections were fixed in 2% Paraformaldehyde (Fluka) for 20 minutes at room temperature, rinsed with PBS and permeabilized with 0.4% Triton X-100 in PBS for 5 minutes at room temperature in Coplin jars. The tissue sections were blocked using Blocking Buffer (2.5% BSA, 0.05% Tween 20 and 10% FCS) for 20 minutes at room temperature inside a humidified chamber. The tissue sections were then incubated with anti-firefly luciferase (Abcam 185924) diluted 1:100 and either anti-CD31 (BD Pharmingen 553370) diluted 1:200 or anti-alpha smooth muscle actin (αSMA) (Abcam Ab7817) diluted 1:100 in Blocking Buffer overnight at 4°C in a humidified chamber. The tissue sections were then washed 3 x 5 minutes in Wash Buffer (PBS containing 0.2% BSA, 0.05% Tween 20) and then incubated with Alexa Flour-488 conjugated secondary antibody (Invitrogen 1874771) diluted 1:400 in Blocking Buffer for 1 hour at room temperature in a humidified chamber. Following 2 x 5-minute washes in Wash Buffer and 1 x 5-minute wash in PBS, the sections were mounted in Vectorshield anti-fade mounting medium containing DAPI (Vector Laboratories). The tissue sections were imaged on a Leica Stellaris 5 confocal microscope, 63x objective, using LAS-X software.

### Haematoxylin and Eosin Staining

Surgically dissected lung tissue was fixed in 10% neutral buffered formalin solution for 24 hours and transferred to 70% Ethanol prior to be processed using Sakura Tissue-Tek VIP® 6 automated tissue processor. Briefly, lung tissues in embedding cassettes were dehydrated by progressing through steps of 70% ethanol for 45 minutes at 37°C, 80% ethanol for 45 minutes at 37°C, 90% ethanol for 30 minutes at 37°C, 96% ethanol for 45 minutes at 37°C, 100% ethanol for 30 minutes at 37°C, 100% ethanol for 1 hour at 37°C, 100% ethanol for 1 hour at 37°C. Dehydrated samples are then cleared by three washes in Xylene for 30 minutes, 45 minutes and 1 hour at 37°C. Finally, specimens are infiltrated by two immersions in 62°C paraffin wax for 45 minutes and 1 hour, followed by two immersions in 62°C paraffin wax for 30 minutes. The tissues were then embedded in paraffin-block using (Leica EG1160 Embedding Center) and 4 μm sections made using ThermoFisher scientific Microtome Microm HM355S and attached to slides.

Prior to staining, sections were deparaffinised by washing slides 3X in HistoclearTM for 2 minutes each, followed by 3 washes 2 minutes each of 100% ethanol, before a final wash 2 min in dH2O. Slides were incubated for 60 seconds in Modified Mayer’s Haematoxylin (Lillie’s Modification) (DAKO), washed for 5 minutes in tap water and immersed for 2 seconds in Eosin followed by washing in dH2O. Prior to mounting coverslips with DPX mounting medium (Sigma) slides were dehydrated by three washes 100% ethanol for 2 minutes each and three washes in Histoclear for 2 minutes each.

The tissue sections were imaged using a Leica DM6000 microscope (10x objective) with a DFC 450 C4 colour camera and Leica LAS-X software.

### Statistical analyses

Microsoft Excel was used for calculations with raw data and GraphPad Prism (version 8) was used for graph generation and statistical analysis. Graphs show the mean of experimental replicates and standard error (SEM), with specific details provided in the figure legends. Multi-group comparisons were tested using one-way ANOVAs with Dunnett’s or Sidak’s correction for multiple comparisons. Pair-wise comparisons were tested using a paired t-test with Holm-Sidak multiple comparison testing when multiple comparisons were made. Details are described in the figure legends.

---

**Supplementary Figure 1:**
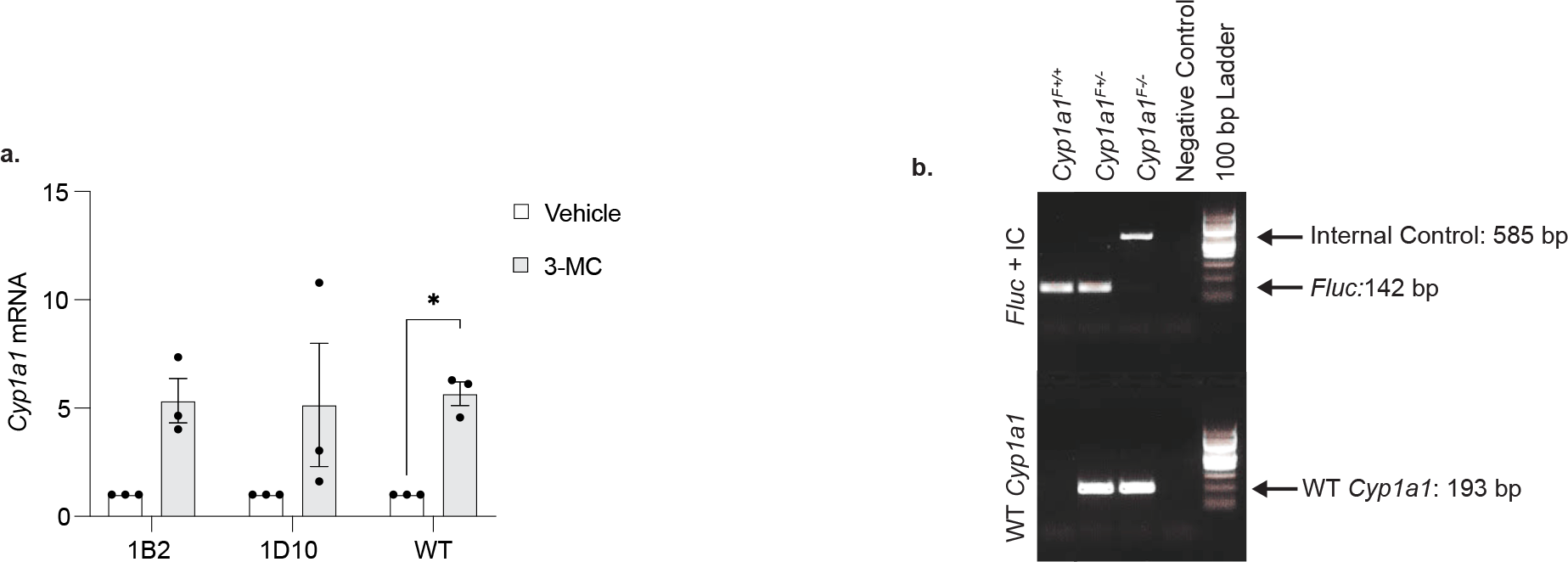
a. RT-qPCR of *Cyp1a1* mRNA expression from two *Cyp1a1^F^* mESC clones (1B2 and 1D10) and WT mESCs following 4-hour 3-MC treatment. Levels of *Cyp1a1* mRNA are normalised to *Gapdh* mRNA and shown relative to the vehicle control. Bars show mean (n=3) +/- SEM with paired t-tests to compare vehicle with 3-MC treated samples. **b.** Representative image of an agarose gel with PCR results for genotyping of *Cyp1a1^F^* mice. Two independent PCR reactions were performed in parallel for each DNA sample. The upper section of the gel illustrates a PCR reaction with two sets of primer pairs: one specific for the *Fluc* gene which amplifies a PCR product of 142 bp and another specific for a region of wild-type *CD79b* (Chr11: 17714036-17714620) that serves as internal control (IC) and amplifies a PCR product of 585 bp. The lower section of the gel illustrates a PCR reaction with only one pair of primers specific for the wild type allele of *Cyp1a1* (WT *Cyp1a1*) and amplifies a PCR product of 193 bp. Arrows indicate the respective sizes of PCR products amplified in the reactions. Examples of homozygous (*Cyp1a1^F+/+^*) and heterozygous (*Cyp1a1^F+/-^*) DNA samples are shown. DNA from the parental Bruce 4 mESC line was used as wild type (*Cyp1a1^F-/-^)* control and the negative control was a non-template PCR reaction. Primers for *Fluc* are labelled as F1 and R1 while primers for WT *Cyp1a1* are labelled as F2 and R2, and their respective sequence locations are indicated in Figure 1a.

**Supplementary Figure 2:**
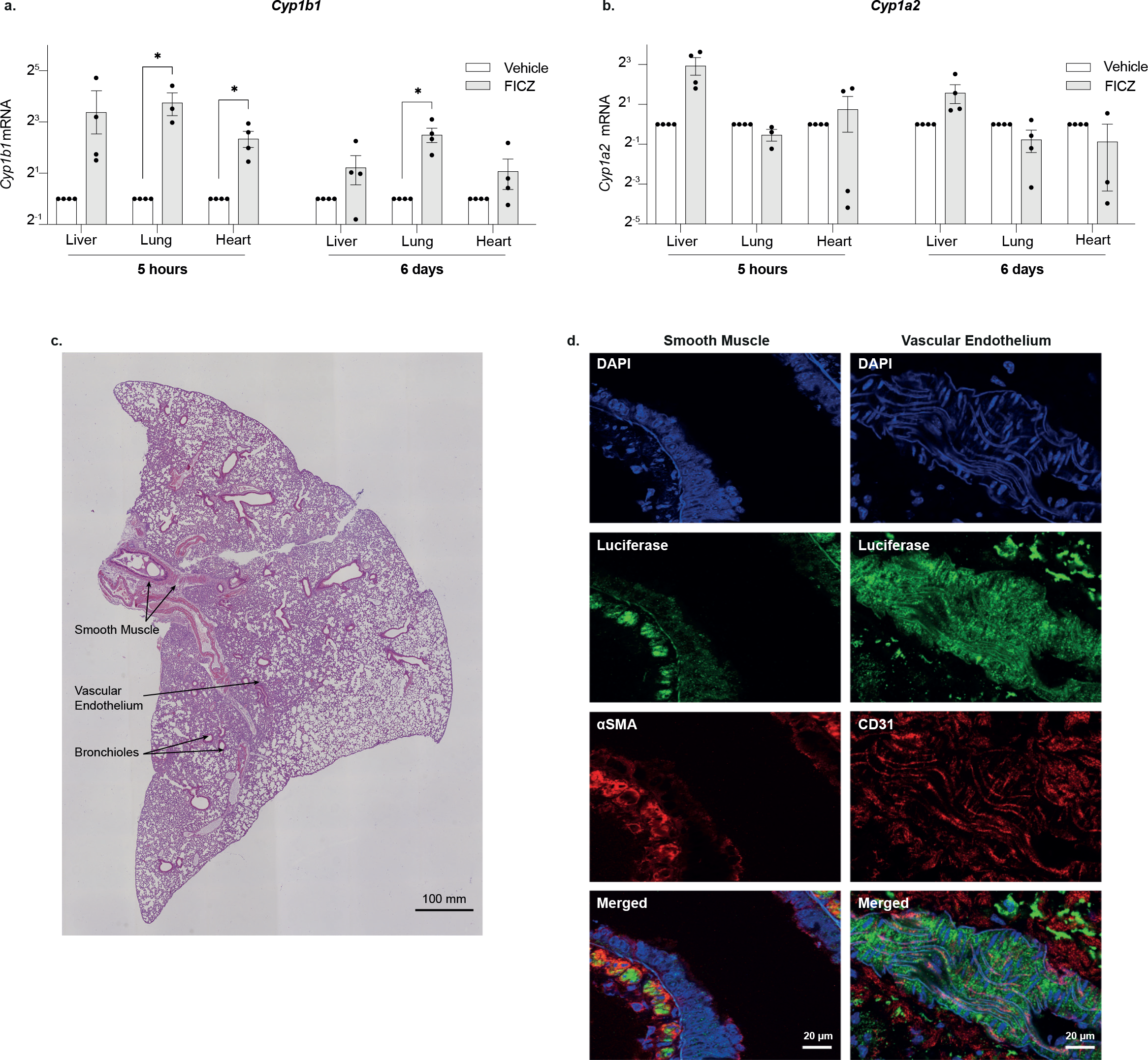
Expression of AHR target genes in the lung following FICZ exposure. **a- b.** Tissues were dissected from adult mice 5 hours and 6 days post FICZ or vehicle injection. RT-qPCR for *Cyp1b1* (**a**) and *Cyp1a2* (**b**). Levels of mRNA were normalised to *18S* rRNA and *Tbp* mRNA and results are shown relative to the corresponding vehicle treated sample. Bars show mean +/- SEM with t-tests with Holm-Sidak multiple comparison testing to compare vehicle with FICZ treated tissues, adjusted p values are shown (*p<0.05). **c**. Adult murine lung tissue 6 days post FICZ treatment was formalin fixed, wax embedded, sectioned at 4 µm and Hematoxylin and Eosin (H&E) stained. Arrows highlight a subset of smooth muscle cells, vascular endothelium and bronchioles identified as luciferase positive by immunofluorescence labelling. **d.** Adult murine lung tissue 6 days post FICZ treatment were cryosectioned at 20 µm and labelled with antibodies characterizing smooth muscle (anti alpha smooth muscle actin, aSMA) or vascular endothelium (CD31), co-detected with anti-firefly luciferase. The left-hand panel shows luciferase positive smooth muscle cells (green) co- stained with aSMA (red), with nuclei counterstained with DAPI (blue) with a merged image below. The right-hand panel shows luciferase positive vascular endothelium (green) co- stained with anti CD31 (red), with nuclei counterstained with DAPI (blue) with a merged image below. Size bars are 20 µm.

**Supplementary Figure 3:**
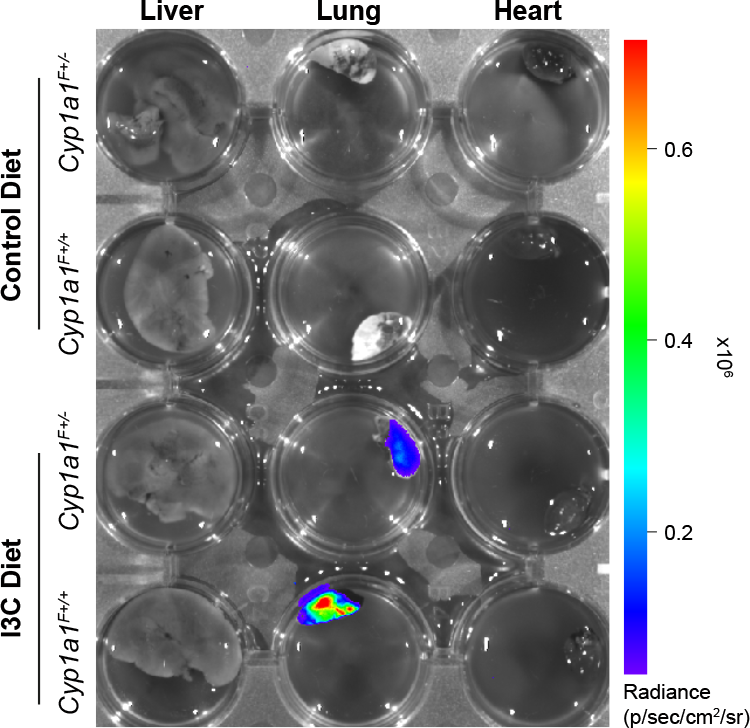
AHR activity in the lung of *Cyp1a1^F^* reporter mice following I3C diet. Corresponding to Figure 4. *Cyp1a1^F+/-^* and *Cyp1a1^F+/+^* adult mice were fed purified control diet or I3C diet for one week following which liver, lung and heart were dissected and analysed by *ex vivo* bioluminescence imaging.

